# Distinctive interaction between cognitive networks and the visual cortex in early blind individuals

**DOI:** 10.1101/437988

**Authors:** Sami Abboud, Laurent Cohen

## Abstract

In early blind individuals, brain activation by a variety of non-perceptual cognitive tasks extends to the visual cortex, while in the sighted it is restricted to supramodal association areas. We hypothesized that such activation results from the integration of different sectors of the visual cortex into typical task-dependent networks. We tested this hypothesis with fMRI in blind and sighted subjects using tasks assessing speech comprehension, incidental long-term memory and both verbal and non-verbal executive control, in addition to collecting resting-state data. All tasks activated the visual cortex in blind relative to sighted subjects, which enabled its segmentation according to task sensitivity. We then assessed the unique brain-scale functional connectivity of the segmented areas during resting state. Language-related seeds were preferentially connected to frontal and temporal language areas; the seed derived from the executive task was connected to the right dorsal frontoparietal executive network; the memory-related seed was uniquely connected to mesial frontoparietal areas involved in episodic memory retrieval. Thus, using a broad set of language, executive, and memory tasks in the same subjects, combined with resting state connectivity, we demonstrate the selective integration of different patches of the visual cortex into brain-scale networks with distinct localization, lateralization, and functional roles.

## 1 Introduction

About one third of the human cortex is commonly labelled as visual. However, imaging studies in sighted humans have shown that some of those regions also carry out multisensory computations and high-order perceptual functions from non-visual input, which persist in early blind individuals (EB; Ricciardi et al. 2014; Murray et al. 2016). In addition, there is functional imaging evidence that in EB, the visual cortex may acquire a novel role in supporting non-perceptual functions (Cecchetti et al. 2016). The scope of such non-perceptual reorganization is large and includes language (Sadato et al. 1996; Amedi et al. 2003; Bedny et al. 2011), mathematics (Kanjlia et al. 2016), memory (Raz et al. 2005; Burton et al. 2012) in addition to activations that may relate to executive functions (Burton et al. 2010; Lewis et al. 2010; Deen et al. 2015). Those activations are unlikely to be epiphenomenal as several studies showed that they correlate with performance measures across participants (Amedi et al. 2003; Lane et al. 2015). In agreement with this interpretation, transcranial magnetic stimulation and lesion studies showed that disrupting the activated areas leads to cognitive impairments in blind subjects (Cohen et al. 1997; Hamilton et al. 2000; Amedi et al. 2004). Moreover, resting-state correlations of the visual cortex with the rest of the brain, often interpreted as resting-state functional connectivity (rsFC), differs between blind and sighted individuals. There is, generally, a decrease in inter-hemispheric occipital rsFC, a decrease in visual to auditory and somatosensory rsFC, and an increase in rsFC between the visual cortex and the lateral prefrontal, parietal and temporal cortices (Liu et al. 2007; Yu et al. 2008; Bedny et al. 2011; Watkins et al. 2012; Butt et al. 2013; Qin et al. 2013; Burton et al. 2014; Wang et al. 2014; Deen et al. 2015; Striem-Amit et al. 2015; see however, Heine et al., 2015).

How are those alterations in resting-state functional connectivity related to the novel cognitive functions acquired by the visual cortex of blind individuals? The goal of this study is to determine whether areas activated by non-perceptual functions in the visual cortex of the EB show functionally relevant brain-scale connectivity.

Addressing a similar question, Kanjlia et al. (2016) showed that distinct visual regions are activated by language and mathematics in early blind individuals. Those two regions were also preferentially connected to distant areas processing language and mathematics, respectively.

In the present study, we assess whether this phenomenon is a more general feature of the visual cortex of the EB. To this end, we use a diverse series of high-level non-perceptual cognitive tasks combined with functional connectivity measures: A novel task switching paradigm directly assessing executive control, and the more customary tasks of speech perception focusing on language comprehension, incidental memory targeting retrieval from long-term memory and a more composite word generation task assessing verbal executive functions. The objective of using both novel and more customary paradigms is to portray a more elaborate description of the functional recruitment of the visual cortex in blind individuals.

Our first prediction, based on studies of memory, language and executive function reviewed above, was that all high-level cognitive tasks should elicit activations in the visual cortex of blind subjects, as compared to sighted controls. In principle, such activations may or may not occupy distinct cortical territories across tasks. Our second prediction, based on Kanjila et al. (2016), was that areas activated by different tasks should be connected to distinct functional networks reflecting the novel involvement of the visual cortex in language, memory and executive functions. If supported, this second prediction would imply that areas activated by different tasks are not entirely identical. In a first step, we studied the activation elicited during the different tasks. In a second step, we used resting-state data in the same subjects, to identify the brain-scale functional networks specifically connected to the activated areas.

## 2 Materials and methods

### 2.1 Subjects

The study included 12 early blind subjects (age: 44±12, mean±SD; blindness onset age: 0) and 16 sighted control subjects (age: 42±12), out of which 12 were matched in sex and age to the group of blind individuals. All subjects were native French speakers. None of the blind subjects had any form of light perception (Table 1). One blind subject was excluded from the analysis due to anomalies in brain anatomy. All subjects signed an informed consent form, were paid for their participation, and were naïve about the aims of the study. The experiment was approved by the local ethical committee.

**Table 1.** Causes of blindness.

| Subject code | Sex | Age | Cause of blindness |
| --- | --- | --- | --- |
| **B1** | M | 25 | Congenital bilateral glaucoma |
| **B2** | F | 28 | Microphthalmia |
| **B3** | F | 54 | Bilateral retinopathy of prematurity |
| **B4** | M | 48 | Retinal detachment |
| **B5** | F | 43 | Microphthalmia |
| **B6** | M | 66 | Congenital retinis pigmentosa |
| **B7** | M | 51 | Unknown |
| **B8** | F | 44 | Bilateral retinopathy of prematurity |
| **B9** | F | 44 | Bilateral retinopathy of prematurity |
| **B10** | M | 44 | Unknown |
| **B11** | F | 31 | Congenital bilateral glaucoma |

### 2.2 fMRI experimental paradigms

Each subject went through 2 magnetic resonance imaging (MRI) sessions performed on two separate days. Before acquisition, subjects participated in short training sessions outside the magnet, during which they performed a few training blocks of the upcoming paradigms. All subjects were blindfolded throughout the MRI acquisitions and were instructed to keep their eyes closed.

#### 2.2.1 Cognitive activation experiments

##### 2.2.1.1 Speech perception

In order to localize the brain regions involved in processing semantics and syntax, we designed a speech perception paradigm in which subjects attended to words and pseudo-words arranged either in sentences or in lists. The 4 types of stimuli were built as follows. Sentences consisted of 96 French sentences ranging from 9 to 12 words (e.g. “Il compte les mots qui couvrent la page du journal” which translates to “He counts the words which cover the page of the newspaper”). Word lists were derived from the sentences by shuffling the words (e.g. “La compte les du Il journal page couvrent qui mots” which translates to “The count the of he newspaper page cover which words”). So-called Jabberwocky stimuli were derived from the sentences by keeping intact the original order of words as well as grammatical morphemes, while replacing the other words with phonologically matched pseudo-words, respecting the original sentence prosody (e.g. “Il sige les fleux qui dasent la plite du gornion”). Pseudo-word lists were derived from the jabberwocky condition by shuffling (e.g. “La sige fleux du plite dasent qui les gornion il”). The 384 (96 × 4) stimuli were recorded by a male native French speaker, and processed with Audacity (v2.1.2, www.audacityteam.org). First, noise removal and normalization (maximum amplitude set to −1.0dB) were applied to the recordings. All stimuli were then duplicated and a pitch change (10% upwards, without changing speed) was applied to the duplicated set in order to generate the same stimuli with a higher-pitch voice. High-pitch stimuli were used only as targets in a difficult voice change detection task aiming at maintaining the subjects’ attention to the stimuli. In order to avoid repetition effects between a sentence and the corresponding shuffled list, stimuli were then split into two sets of 48 stimuli per category (sets A and B). Two groups of stimuli were then created: a group consisting of sentences from set A and lists from set B (group 1) and conversely (group 2). Subjects were randomly assigned to one of the two groups.

Subjects received 16 blocks of each of the 4 categories. Blocks with a duration of 12 s were arranged such that each block contained 3 stimuli and an inter-stimulus silence of 0.6 s. For each condition, 3 blocks out of 16 were selected at random in which one stimulus was substituted by its high-pitched counterpart. Subjects had to press the left button as soon as they detected this voice change. Due to the varying duration of the stimuli and in order to form blocks homogenous in duration, we adjusted the duration of stimuli in the range of −5% to 7%, without changing their pitch. In order to equate the duration of blocks, we grouped stimuli into blocks using an optimization algorithm (constrained linear least-squares solver) for each condition. A pseudo random order of the blocks was generated for each subject. No more than 2 consecutive blocks of the same condition were allowed. Each block ended with a rest period (2 s), and 24 blocks of rest (10 s) were randomly distributed among the task blocks. The experiment was divided into 2 equal runs (9 m 43 s each) to minimize fatigue and movement. Each run started and ended with additional rest periods (20 s and 10 s, respectively).

Blocks were tagged as erroneous if the subject either failed to detect the change in the speaker voice, or signaled a change when there was none. The effects of semantics were assessed using a contrast of (Sentences + Word-lists) minus (Jabberwocky + Pseudoword lists). The effects of syntax were assessed using a contrast of (Sentences + Jabberwocky) minus (Word-lists + Pseudoword-lists).

##### 2.2.1.2 Word generation

In order to probe verbal executive processing, we used a word generation paradigm. Subjects were presented with auditory nouns, and had to overtly generate words according to 3 tasks: Repeat, where subjects had to repeat the heard noun (e.g. cheese -> cheese), Verb, where subjects had to generate a verb associated to the heard noun (e.g. cheese -> to eat), and Initial, where subjects had to generate a word starting with the same phoneme as the heard noun (e.g. cheese -> children).

Stimuli consisted of a set of 70 French nouns selected from the Lexique database (New et al. 2004). Instructions consisted of the words “répéter”, “verbe”, and “initiale” which translate to “repeat”, “verb” and “initial”, respectively. Stimuli and instructions were synthetically generated using the Text-to-Speech function built in OSx 10.9.5 (Apple Inc., CA, United States). Subjects received 14 stimulation blocks for each of the 3 conditions. Each block started with an instruction word (2 s), followed by 5 trials (3 s each) including stimulus presentation and response time-window, and ended with a resting period (3 s), totaling 20 s per block. Pseudo-random permutations of stimuli and block types were generated for each subject. No more than 2 consecutive blocks of the same condition were allowed, and subjects received all 70 words under the 3 conditions. In addition, 18 rest blocks (12 s) were randomly distributed among the stimulation blocks. The experiment was divided into 2 equal runs (9 min 6s each) to minimize subject fatigue and movement. Each run started and ended with an additional rest period (20s and 10s, respectively). Subjects were instructed to produce an overt response after each trial, speaking with a normal voice while maintaining their head still.

Responses were recorded for subsequent scoring of errors. Recordings were denoised so as to remove the scanner noise, and latencies were computed as the lag between the onset of stimuli and the onset of vocal responses. This lag was verified and, when needed, corrected manually for all trials. Spurious preverbal noises were manually excluded from the computation of latencies. Trials on which subjects either did not respond in the designated time-window or produced a response that did not comply with the current instruction were flagged as errors and excluded from the response time analyses. One blind subject with excessive head motion and two sighted subjects tested with different scanning parameters were excluded from the analysis of this experiment. The effects of word generation were assessed using the Verb minus Repeat and Initial minus Repeat contrasts. We hypothesized that the Initial task would be more effortful than the Verb task due to the need for an active lexical search when compared with the Verb task which is more associative in nature, thus, resulting in a differential brain response.

##### 2.2.1.3 Executive control

We used an auditory adaption of a contextual task-switching paradigm (Koechlin et al. 2003). This paradigm is thought to be sensitive to task-set reconfiguration costs, the process of manipulating stimulus-response rules (Monsell 2003). Trials consisted of one French vowel ([a],[i], or [u]) followed by a series of 3 or 4 different notes played on the piano, forming an ascending or a descending scale. On each trial, the initial vowel specified the task to be performed. For vowel [a], the task was to judge whether the upcoming series contained 3 or 4 notes, and to respond with a left or a right button press, respectively. For vowel [u], the task was to judge whether the scale was descending or ascending, and to respond with a left or a right button press, respectively. Finally, for vowel [i], no response was to be made. The [a], [u], and [i] vowels were recordings (500 ms) of a male human voice, normalized and low-pass filtered (3500 kHz). Piano notes were generated using the Grand Piano instrument in Kontak 4 (Native Instruments GmbH, Germany). The notes were C2, E2, G#2, C3, E3, G#3, C4, and their duration was 125 ms. The full set of stimuli thus included 12 different items: 3 tasks × 2 numbers of notes × 2 melodic contours. Those items were used to build trials for the 3 experimental conditions: Number blocks which included only [a] and [i] trials; Melody blocks which included only [u] and [i] trials; and Mixture blocks which included all 3 possible tasks (i.e. [a], [u], and [i]). Each block started with the synthetically generated instruction word “nombre”, “mélodie”, or “mélange”. Subjects received 8 blocks of Number and 8 blocks of Melody (which will be referred to as Single-task blocks), and 16 blocks of Mixture (which will be referred to as Dual-task blocks). Each block thus began with an auditory instruction (2s), followed by 9 trials (3 s each) during which a vowel and a series of notes were played. Each block ended with 2 s of rest, for a total of 31 s. For each subject, a pseudorandom permutation of block order was generated. No more than 2 consecutive blocks of the same condition were allowed. In addition, a rest block (10 s) preceded each task block. Several constraints were enforced when generating the random order of the stimuli within blocks: 1) A block cannot start with an ‘i’ stimulus (no response), 2) No consecutive ‘i’ stimuli are allowed, 3) No more than 2 consecutive ‘a’ or ‘u’ stimuli are allowed, 4) The number of left and right presses are equal in each block, 5) The number of congruent trials (same button for the Number and Melody tasks) and incongruent trials (different button for the two tasks) are equal in each block. The experiment was divided into 2 equal runs (11 min 1 s each) to minimize fatigue and movement. Each run ended with an additional rest period (10 s). Subjects held a response button in each hand and were instructed to respond as soon as they could without compromising performance. Response time was defined as the lag between stimulus onset and button press. Whenever subjects failed to respond in the designated time window, pressed the wrong button, or responded in the no-response condition (i.e. on trials with a [i] vowel), trials were tagged as errors and excluded from the response time analyses. One blind and one sighted subjects with excessive head motion and two sighted subjects with different scanning parameters were excluded from the analysis of this experiment. The effects of non-verbal executive control were assessed using the Dual-task minus Single-task contrast.

##### 2.2.1.4 Long-term memory

In order to localize regions involved in long-term memory, we used an incidental long-term memory paradigm, in which subjects were asked to judge, when presented with sentences, whether they had previously heard them or not. A variant of the old/new recognition task, it is often used in order to reveal brain regions involved in retrieval from episodic memory (Konishi et al. 2000; Rugg and Henson 2002; Spaniol et al. 2009). There were 2 experimental and 2 baseline conditions in the memory experiment. Experimental conditions: Old sentences that subjects had heard as part of the Sentence condition of the speech perception experiment; New sentences that had never been presented before. Baseline conditions: Instruction sentences asking subjects to press a button, for each one of the two response buttons (e.g. “Merci d’appuyer maintenant sur le bouton de droite” which translates to “Please press now on the right button”). Subjects were not aware of the existence of this experiment beforehand so they would not engage in active encoding during the Speech perception experiment. It was introduced 15 minutes after the speech perception experiment, during which anatomical images were acquired. Using sentences from the speech perception experiment created a double benefit. First, it allowed probing the activation by similar perceptual input under different tasks. Second, it eliminated the need to explicitly teach subjects a new set of stimuli, a time consuming process that would have reduced testing time.

The material comprised 60 sentences taken from the speech perception experiment and 60 new sentences (recorded and processed as previously described). For each old sentence, a new sentence was created by using the structure of the former with changes limited to the content (e.g. “La maison où habite cet homme est très abîmée” which translates to “The house where this man lives is very damaged” became “La forêt où vivent ces oiseaux doit être abattue” which translates to “The forest where those birds live should be cut down”), thus achieving a matching in syntax and number of words (New: 9.8±1.8, Old: 9.9±1.4 (mean±sd); t(59)=-0.75, p=0.43, paired t-test). There were 10 instruction sentences (5 for each button side). Stimuli were divided into two groups following the division in the speech perception experiment, so as to match the specific set of sentences which each subject had heard before. We used a fast event-related design, with 30 sentences for each condition. Each trial had a total duration of 5.5 s, and consisted of a sentence, followed by a GO signal (a short ringing sound; 250 ms) and the response time-window. Subjects were instructed to respond by pressing the right-hand button for old sentences and the left-hand button for new sentences. In the Instruction condition, subjects had to obey the auditory instruction and press accordingly. The Instruction sentences were used as a baseline condition for new and old sentences following the response-button mapping. A pseudorandom order of events was generated for each subject with no more than 2 consecutive trials of the same condition allowed. 27 periods of rest (6 s) were distributed among the task trials. The experiment was conducted in one run (11 min 12 s), which started and ended with additional rest periods (10 s and 5 s, respectively). Trials were classified according to subject response. Old sentences that were correctly detected were classified as hits, old sentences that were identified as new were classified as misses, new sentences that were identified as such were classified as correct rejections and new sentences that were identified as old were classified as false positives. Only correctly classified trials were used for assessing the brain response in this experiment. Data from one blind subject that failed to comply with task instructions and from two sighted subjects that made 8/30 and 15/30 mistakes in the Instruction conditions were excluded from the analysis of this experiment. The general effects of the memory task were assessed using the Hits minus Instruction contrast. The effects of retrieval from long-term memory were assessed using the Hits minus Correct rejections contrast after subtraction of their respective Instruction conditions.

#### 2.2.2 Experimental setup

##### 2.2.2.1 Stimulation methods

We used Psychophysics Toolbox Version 3 (Brainard 1997; Pelli 1997; Kleiner et al. 2007) for MATLAB (Release 2011a The MathWorks, Inc., Massachusetts, United States) to implement the experimental design. The auditory stimuli were relayed to an MR Confon Easy 04 amplifier which was connected to MR Confon HP SI 01 headphones (MR confon GmbH, Germany). Subject vocal responses were recorded using either an Optoacoustics FOMRI III microphone (Optoacoustics Ltd., Israel) or a Sennheiser MO 2000 set (Sennheiser Electronic GmbH & Co., Germany). The microphone was attached to the head coil and was placed at an approximate distance of 1 cm from the subject’s mouth. All subjects wore foam earplugs and blindfolds prior to installation in the scanner.

##### 2.2.2.2 Testing sessions

The first session included (i) a T1 anatomical volume, (ii) fMRI images during resting state, (iii) the two runs of the executive control experiment, and (iv) the two runs of the word generation experiment. The second session included (i) the two runs of the speech perception experiment, (ii) T1 and T2 anatomical volumes, (iii) the long-term memory experiment.

#### 2.2.3 Behavioral data analysis

In the word generation and executive control experiments, median latencies and individual error rates were entered into ANOVAs with task as within-subject factor, group as between-subject factor and subjects as random factor. Post-hoc Tukey’s tests were used to compute differences between conditions in each group. In the speech perception experiment, unpaired t-tests were used to compare the groups on error rates. In the long-term memory, one-sample t-tests were used to compare d’ values to chance and an unpaired t-test was used to compare the d’ values between the groups.

### 2.3 fMRI acquisition and analysis

#### 2.3.1 Imaging parameters

Anatomical and functional brain images were acquired using a 3-tesla MRI (Verio, Siemens AG, Germany). Using a 32-channel head coil, we acquired a three-dimensional T1-weighted anatomical volume (MP RAGE sequence, TR=2300ms, TE=3.1ms, flip angle=9°, 0.8mm iso-voxel resolution) and a 10 minutes series of whole-brain resting-state BOLD sensitive images (gradient-echo (GE) echo planar imaging (EPI) sequence, 45 slices, slice thickness/gap =3/0.6mm, FOV 192 × 192mm, A>>P phase encoding direction, TR=2500ms, TE=30ms, flip angle=80°). For all task-related activation experiments, we used a 12-channel head coil, to acquire a whole-brain BOLD sensitive contrast (GE-EPI sequence, 40 slices, slice thickness/gap=3/0.3mm, FOV 204 × 204mm, A>>P phase encoding direction, TR=2020ms, TE=25ms, flip angle=80°).

#### 2.3.2 Data analysis

##### 2.3.2.1 Preprocessing

Preprocessing was performed using SPM12 (http://www.fil.ion.ucl.ac.uk/spm/) and MATLAB (Release 2014b). Anatomical volumes were segmented and normalized to the standard Montreal Neurological Institute (MNI) stereotactic space. All functional time-series were slice-time corrected, motion corrected to the mean functional imaging using a tri-linear interpolation with six degrees of freedom, co-registered to the anatomical volume of the first session, normalized, resampled (to 3mm cubic voxels), and spatially smoothed (5mm FWHM, isotropic). Resting-state functional volumes were further preprocessed using CONN toolbox (Whitfield-Gabrieli and Nieto-Castanon 2012) in order to bring spurious correlations to a minimum. We applied linear trend removal to correct for signal drift and band-pass filtering (0.008-0.09Hz) to reduce non-neuronal contributions to the spontaneous BOLD fluctuations. Moreover, we used the aCompCor strategy (Behzadi et al. 2007) in addition to the estimated subject motion parameters (6 rigid-body head motion parameter values – x, y, z translations and rotations) to define temporal confounding factors due to head motion. Finally, we compared the maximum absolute head motion value between the groups of sighted and blind subjects and found no significant difference (t(22)=0.43, p=0.67, unpaired t-test; Van Dijk et al. 2012)

##### 2.3.2.2 Analyses of task-related activation paradigms

For the single-subject analyses, we used general linear-models (GLM) with paradigm specific regressors, in addition to 6 motion parameters and a high-pass filter (128 s cutoff, except in the case of the word generation paradigm, where it was set to 330 s). A boxcar function using the onsets and durations of the following predictors was convolved with a canonical hemodynamic response function (HRF) to generate the GLM regressors. Speech perception experiment: one regressor for each of the 4 conditions, plus one regressor for target sentences and one for button presses. Word generation experiment: one regressor for each of the 3 conditions plus one regressor for instructions. Executive control experiment: one regressor for each of the 3 conditions plus one regressor for instructions. Long-term memory experiment: one regressor per response class, i.e., hits, misses, correct rejections and false positives, one regressor per instruction condition, i.e., right and left plus two regressors for left and right button presses. For second-level group analyses we used one-sample t-tests for inferences on a single group of subjects, and two sample t-tests to assess differences or conjunctions involving the two groups. We used the following procedure to correct for multiple comparisons on the cluster-level:

First, we set the second-level voxel-wise cluster-forming threshold to p<0.001 for the word generation and speech perception experiments, and p<0.005 for the long-term memory and executive control experiments in order to increase sensitivity. Then, the square-root of the residuals image of the GLM result of each subject was computed using 3dcalc (AFNI). Image smoothness was estimated using 3dFWHMx (AFNI) for each subject, assuming it follows a mixed autocorrelation function (ACF) and not a simple Gaussian ACF (Cox et al. 2017). The average smoothness of the group under investigation (sighted, blind, or sighted and blind together) was then used in a Monte Carlo simulation (10,000 iterations; 3dClustSim), to compute a minimal cluster size k for each second-level result. This k value calculation corresponded respectively to k=25, 23, 25 in the sighted, the blind and both groups together in speech perception; k=24, 25, 23 in word generation; k=80, 92 and 87 in executive control; and k=80, 73, 76 in long-term memory. Thus, all the reported activation maps show significant results that are cluster-wise corrected (p<0.05).

##### 2.3.2.3 Functional connectivity analysis

One representative contrast from each task-related activation experiment was used in a functional connectivity (FC) analysis on the resting-state data (Fox and Raichle 2007). Seed regions were defined by intersecting the activation map (in blind subjects) of each representative contrast with a mask of the visual cortex. The mask included the gray-matter voxels of bilateral hOc1-2, hOc3v/d, hOc4v/d/la/lp, hO5 and FG1-4 (Anatomy Toolbox, Eickhoff et al. 2005) in addition to the middle and superior occipital gyri (AAL, Tzourio-Mazoyer et al. 2002). Activation maps were cluster-wise corrected (p<0.05) across the whole brain following voxel-wise cluster-forming thresholding of p<0.001 to achieve restricted seed regions. In the Executive Control contrast, a cluster-forming threshold of p<0.001 was too strict leading to the use of a more sensitive threshold of p<0.005.

Moreover, all activated voxels constituting the seed regions are also significant after voxel-wise FDR correction for multiple comparisons (q=0.05) within the visual cortex mask (Fig. 2). FC was computed as follows. For each subject, the average time-course of the preprocessed resting-state BOLD signal was extracted from each seed region, and a whole brain correlation map was computed. Correlation values were then transformed into Z values (Fisher’s r-to-Z). For each seed, the individual Z-maps were entered in a second-level group analysis where the groups of blind and sighted subjects were compared resulting in a t-map of connectivity differences (CONN toolbox). Next, in order to generate group-level functional connectivity maps representing the unique connectivity of each seed, we used semi-partial correlation coefficients. For each seed, those coefficients result from a correlation analysis of its time-course while regressing out the time-courses of all other seeds. In other words, semi-partial correlations depend only on seed-specific variance. It should be noted here that this method does not impose constraints on the spatial extent of the resulting long-distance connectivity. Thus, different seeds can be connected to the same target regions, due to unique variance that distinguishes them from one another. In all resting-state connectivity analyses we used a voxel-wise cluster-forming threshold of p<0.005 and a cluster-wise threshold of p<0.05 (FDR corrected).

**Figure 1.**
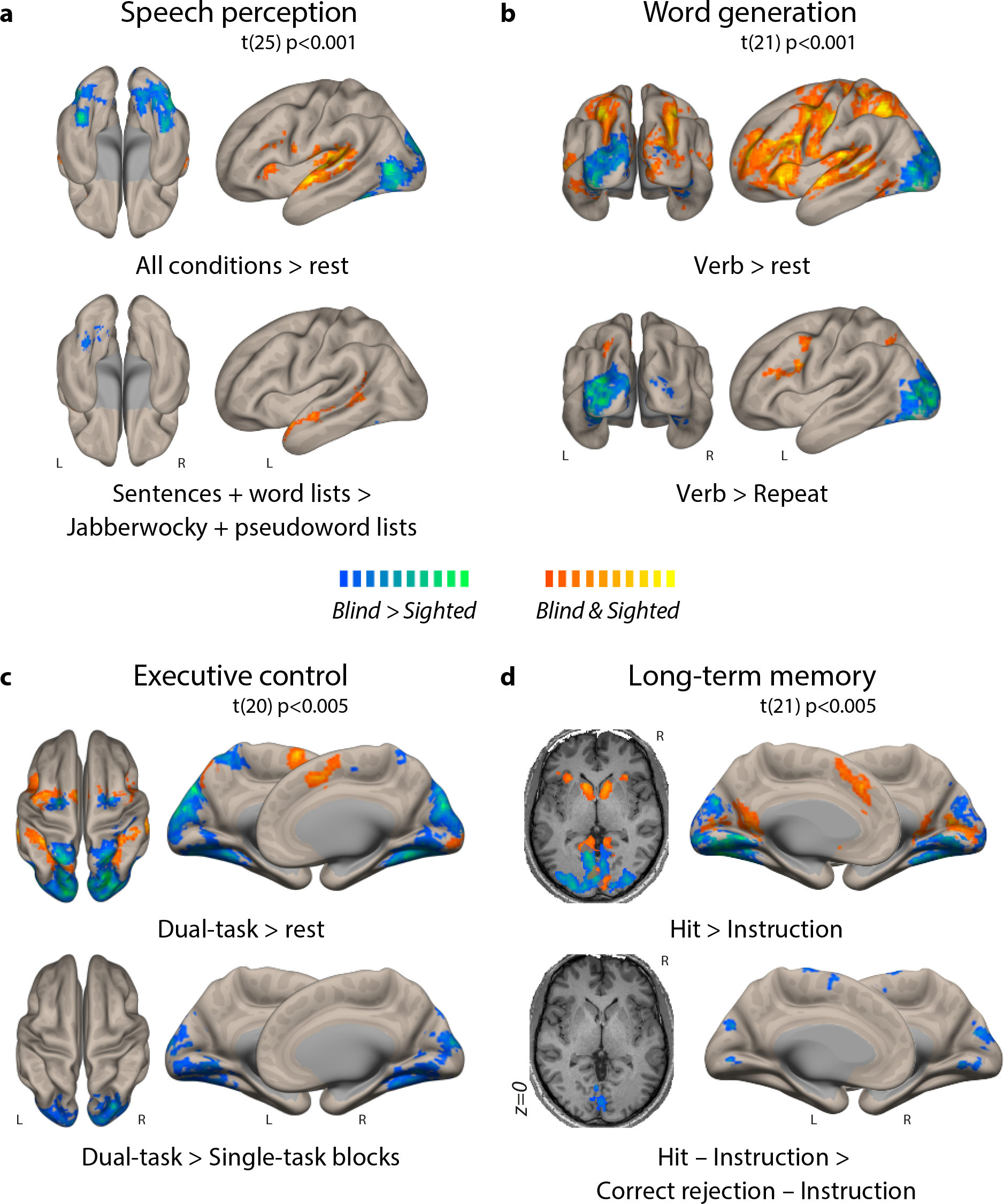
Activation in blind and sighted subjects (conjunction and differences) Whole-brain activation of selected conditions minus rest, and minus high-level control in the (**a**) speech perception, (**b**) word generation, (**c**) executive control and (**d**) long-term memory experiments. For each contrast, hot colors show significant activation in the conjunction of blind and sighted subjects, and cold colors show activation significantly stronger in blind than in sighted subjects. There was no activation stronger in the sighted than in the blind. With contrasts using high-level control conditions, all experiments activated the visual cortex in the blind relative to the sighted. Voxelwise cluster-forming thresholds are reported in the figure, and the upper end of the color bar range was set to p<1.0e-07. All maps are clusterwise corrected (p<0.05).

**Figure 2.**
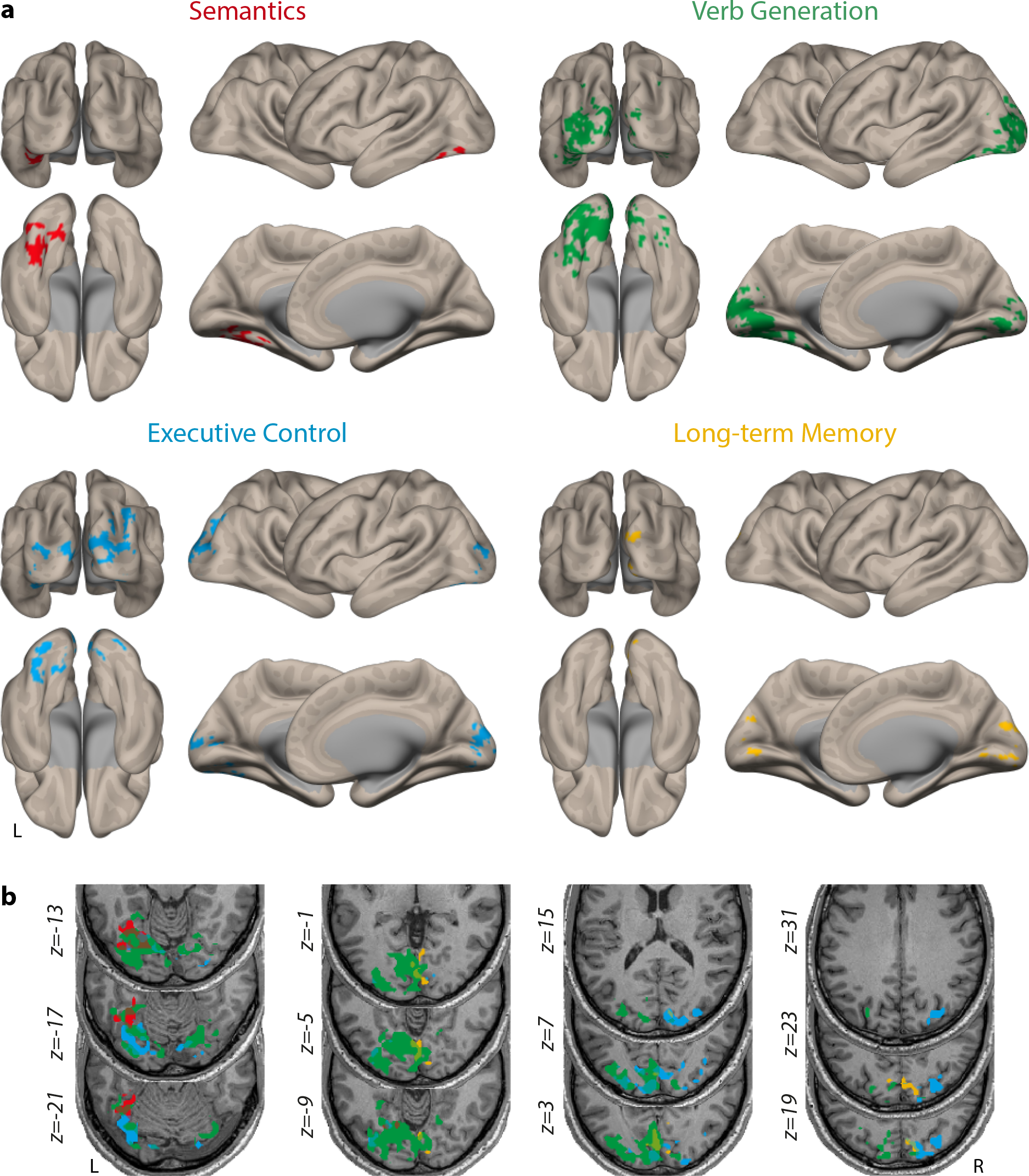
Occipital seeds for the functional connectivity study. Surface (**a**) and slice (**b**) views of the activation by the four representative contrasts, in the visual cortex of the blind. Semantics in shown in red (contrast: [Sentences + Word-lists] minus [Jabberwocky + Pseudo-word lists]), Verb Generation in green (contrast: Verb generation minus Word repetition), Executive Control in blue (contrast: Dual-task minus Single-task), and Long-term Memory in yellow (contrast: Hits minus Correct rejections, with subtraction of the baseline condition). Those maps were used as seed regions in the functional connectivity analysis.

##### 2.3.2.4 Task-related activation in connected ROIs

To test whether the uniquely connected regions found in the previous analysis are sensitive to the task-related contrasts that were used to define the seeds that yielded them, we devised the following analysis in individual subjects. First, for each group-level unique connectivity map, we defined individual-subject ROIs in the connected long-distance regions. For connected regions that encompassed different cortical zones, such as the prefrontal and temporal cortices in the unique connectivity from the Semantics seed, we defined an ROI per zone. The zones were defined using the AAL atlas for each of the four contrasts, for a total of seven zones (Table 3, 2^nd^ and 3^rd^ columns). For the Semantics seed, two zones were defined: 1) left prefrontal (Left Frontal Pole, Superior Frontal Gyrus, Middle Frontal Gyrus, Inferior Frontal Gyrus (pars triangularis and pars opercularis)); and 2) left superior/middle temporal (Left Superior Temporal Gyrus (anterior and posterior divisions), Middle Temporal Gyrus (anterior & posterior divisions, and temporooccipital part)). For Verb generation, we defined a left prefrontal zone (equivalent to region 1 in Semantics). For executive control, two zones were defined: 1) right prefrontal (symmetrical to prefrontal regions used in Semantics); and 2) right superior parietal (Right Superior Parietal Lobule, Lateral Occipital Cortex (superior division)). Finally, for long-term memory, also two zones were defined: 1) anterior cingulate (Frontal Medial Cortex, Cingulate Gyrus (anterior devision), and Right & Left Paracingulate Gyrus); 2) precuneus (Precuneus Cortex while removing voxels under MNI coordinate z=35 to exclude visual cortex). Then, ROIs were defined as the intersection of those anatomical zones with the corresponding individual unique connectivity map (thresholded at p<0.01 uncorrected). This resulted in 7*9 ROIs (zones * subjects). Second, we extracted the average value of the contrast of interest from each ROI for each subject. For example, from the individual-subject ROIs defined in the prefrontal zone from the verb generation connectivity maps, we extracted the value of the Verb minus Repeat contrast. Third, we used one-sample t-tests across subjects to assess whether those ROIs were significantly activated by the corresponding contrasts of interest (FDR-corrected). In this analysis, we only used subjects that had data in all experiments. See also section 1.4 in supplementary material for testing all contrasts in all ROIs.

##### 2.3.2.5 Variance explained in connected ROIs

Using the same ROIs defined above, we computed the percent variance uniquely explained by the corresponding seed, by squaring the semi-partial correlation coefficient in each voxel and averaging across voxels, and then across blind subjects. Moreover, we computed for the same ROIs the percent variance explained by combining all seeds together. To this end, we computed for each voxel a regression model with the time-course of the four seeds as independent variables. The resulting R^2^ from this regression was averaged across voxels and then across subjects.

### 2.4 Display

Slice views were generated using BrainVoyager QX (2.8.4, Brain Innovation, The Netherlands) from thresholded SPMs converted to VMP format using Neuroelf (1.0, http://neuroelf.net/) and overlaid on an MNI normalized anatomy converted to VMR format. Surface views were generated using CONN toolbox Surface display, which projects thresholded SPMs to a FreeSurfer surface registered to MNI space.

## 3 Results

### 3.1 Behavioral results

We first quantified the performance of the blind and sighted subjects in all tasks.

#### 3.1.1 Speech perception

The rate of miss (43.4%) and false-positive (6.5%) errors did not differ between blind and sighted subjects (t(26)=0.28, p=0.78, and t(26)=−0.19, p=0.85, unpaired t-test).

#### 3.1.2 Word generation

There was a significant effect of task (F(2,42)=315.15, p<1.0e−5, ANOVA). As expected, the Verb (1557 ms) and Initial (1840 ms) conditions were slower than the Repeat condition (1041 ms); and the Initial was slower than the Verb condition (p<1.0e−4 for all comparisons, Tukey’s test). There was no significant difference between groups (F(1,21)=0.52, p=0.48) and no task × group interaction (F(2,42)=2.45, p=0.10). Error rates followed the same pattern, also showing an effect of task (F(2,42)=31.52, p<1.0e−5). More errors were made in the Verb (8.6%) and Initial (13.6%) conditions than in the Repeat condition (0.4%) and the Initial condition had the most errors (p<1.0e−4 for Verb and Initial minus Repeat and p=0.01 for Initial minus Verb, Tukey’s test). There was no significant difference between groups (F(1,21)=0.1, p=0.76) and no task × group interaction (F(2,42)=0.71, p=0.50).

#### 3.1.3 Executive control

There was a significant effect of task (F(1,20)=33.1, p<1.0e−4) and a task x group interaction (F(1,20)=6.45, p=0.02): The dual task was slower than the single task, and this difference was larger in the blind (mean: Dual 1543 ms, Single 1424 ms, t(9)=4.33, p=0.002, paired t-test) than in the sighted subjects (mean: Dual 1422 ms, Single 1375 ms, t(12)=3.98, p=0.002, paired t-test). There was no overall difference between groups (F(1,20)=2.96, p=0.1, ANOVA). Error rates were low and showed no effect of group (mean: Blind 4%, Sighted 6.4%, F(1,20)=0.81, p=0.38, ANOVA). There was, however, an effect of condition (F(1,20)=8.3, p=0.008, ANOVA) where the Dual-task condition had a higher error rate than the Single-task condition (3.1% and 2.3%, respectively).

#### 3.1.4 Long-term memory

Both groups performed well above chance with a mean d’ value 1.46±0.41 (mean±std, t(9)=11.14, p<1.0e−5, one-sample t-test) in blind and 1.00±0.66 in sighted subjects (t(13)=5.644, p<1.0e−4, one-sample t-test), with a non-significant trend to the advantage of blind subjects (t(22)=1.92, p=0.068, unpaired t-test).

### 3.2 Task-related activations

For each experiment and each contrast, we will present in turn the activations common to sighted and blind subjects, and the activations which are larger in the blind than in the sighted, with an emphasis on visual cortex. Note that we found no activations larger in the sighted than in the blind subjects. The separate results of the two groups may be found as supplemental material and will be referred to where relevant.

#### 3.2.1 Speech perception

In this experiment, subjects performed a non-language task in order to ensure constant attentional engagement between the different conditions of interest. In both groups, the contrast of the four conditions (Sentences, Word-lists, Jabberwocky, Pseudo-word lists) minus rest activated bilateral primary and associative auditory areas in addition to bilateral inferior frontal sulcus (IFS), left inferior frontal gyrus (IFG) and bilateral medial superior frontal cortex (SFC; Fig. 1a). Moreover, in the same contrast, there was stronger activation in bilateral visual cortex of the blind subjects, spanning its lateral, media and ventral aspects (Fig. 1a; Supplementary Fig. 1a for separate group activations).

We assessed the main effect of semantics by contrasting (Sentences + Word-lists) minus (Jabberwocky + Pseudo-word lists) (Fig. 1a; Supplementary Fig. 1a; Supplementary Fig. 2a). There was a left-predominant activation common to both groups along the bilateral superior temporal sulcus (STS). Stronger activation was found in the left anterior fusiform gyrus of the blind subjects, overlapping with cytoarchitectonic areas FG1, 3 and 4 (peak coordinates x=-34, y=-55, z=-14; area nomenclature following the SPM Anatomy toolbox, Eickhoff et al. 2005).

We then tested the main effect of syntax by contrasting (Sentences + Jabberwocky) minus (Word-lists + Pseudo-word lists). In sighted subjects, we found activations predominantly in the left STS and IFG (Supplementary Fig. 1a). In blind subjects, however, the same regions were activated below significance threshold which is why we found no activation common to both groups. Also below significance was an activation in the left lingual gyrus when comparing the blind minus the sighted subjects.

Due to the lack of a significant result in the main effect of syntax, we will only use the main effect of semantics in the following analysis and it will be referred to as the “Semantics” contrast.

#### 3.2.2 Word generation

The contrast of the Verb minus Repeat conditions showed activation common to both groups in the left IFS and left-predominant medial SFC (Fig. 1b). Activations were larger in blind than in sighted subjects in left inferior and lateral occipital cortex and hOc2 on the medial face of the occipital cortex. In the right hemisphere, the differences were significant in the right ventral stream anterior to hOc3v (Fig. 1b; Supplementary Fig. 1b; Supplementary Fig. 2b). The Initial minus Repeat contrast showed a very similar activation pattern to the Verb minus Repeat contrast (Supplementary Fig. 3). The comparisons between Initial and Verb conditions showed neither significant common activation nor significant group differences.

Since the Initial and the Verb conditions did not differ and considering that the Verb condition had been used in previous studies (Burton et al. 2002; Amedi et al. 2003; Ofan and Zohary 2007), it was chosen to represent verbal executive functions in the following analysis and be referred to as the “Verb generation” contrast.

#### 3.2.3 Executive control

The Dual-task minus rest contrast showed bilateral activations common to both groups, encompassing the auditory (primary and associative), prefrontal (IFS, IFG), premotor, and medial SFC cortices in addition to the intra-parietal sulcus (IPS) and the frontal eye-fields (FEF; Fig. 1c).

Activations were stronger in the blind subjects in bilateral inferior, lateral and medial occipital cortex extending to the posterior parietal cortex and to the superior parietal lobule, in addition to a region posterior to the FEF (Fig. 1c; Supplementary Fig. 1c).

The Dual-task minus Single-task contrast showed no activation common to both groups, due to a difference in lateralization of prefrontal activations between groups. Indeed in sighted subjects, the activation was located in the left anterior dorsolateral prefrontal cortex (DLPFC). In the blind, however, we found activations in the right DLPFC and right IPS in addition to a right-predominant occipital activation (Supplementary Fig. 1c). Stronger activation was found bilaterally in the blind subjects in a right-predominant region extending from the lateral occipital cortex to the posterior parietal cortex (peak coordinates x=30, y=-80, z=18), in addition to parts of the right medial ventral stream (hOc4v and FG3), left posterior ventral stream (hOc4v, FG1), left lingual gyrus, and right medial occipital cortex (hOc2; Fig. 1c; Supplementary Fig. 2c).

The Dual-task minus Single-task contrast will be referred to as the “Executive Control” contrast in the following analysis.

#### 3.2.4 Long-term memory

In both groups, a broad contrast sensitive to the memory task (Hit minus Instruction) activated the bilateral posterior cingulate cortex (PCC), heads of caudate, anterior insula, medial SFC, and primary visual cortex (hOc1; Fig. 1d). Stronger activation was found in bilateral inferior, lateral and medial visual occipital cortex in addition to the right anterior IPS of the blind subjects (Fig. 1d; Supplementary Fig. 1d).

We then used a more specific contrast of successful retrieval from episodic memory (Hits minus Correct rejections while subtracting the baseline Instruction condition; see Materials and Methods). This contrast showed no significantly activated clusters surviving the correction for multiple comparisons in the conjunction of both groups. However, clusters activated below extent threshold can be found in the right superior frontal gyrus (SFG), anterior cingulate cortex (ACC), right head of caudate, left angular gyrus and left cuneus (Table 2). All of which were previously shown to be implicated in memory retrieval (Spaniol et al. 2009). In the same contrast, sensitive to successful retrieval, stronger activation in blind subjects was found bilaterally in the medial occipital cortex overlapping with hOc1,2 and hOc3d (peak coordinates x=-2, y=-86, z=4 in left hOc1; Fig. 1d; Supplementary Fig. 1d; Supplementary Fig. 2d).

**Table 2.** Clusters activated below corrected extent threshold for the successful retrieval contrast in the conjunction of the blind and the sighted.

| Cluster location | Cluster size | Peak Z | Peak p(unc) | x,y,z {mm} |
| --- | --- | --- | --- | --- |
| Superior prefrontal | 28 | 3.71 | 0.000 | 48 35 20 |
| Head of caudate | 8 | 3.41 | 0.000 | 9 8 -1 |
| Anterior cingulate cortex | 31 | 3.30 | 0.000 | -3 44 5 |
| Cuneus | 7 | 3.10 | 0.001 | -6 -61 38 |
| Angular | 5 | 3.01 | 0.001 | -45 -55 44 |

The successful retrieval contrast, representing the activation for long-term memory will be referred to as the “Long-term Memory” contrast in the following analysis.

### 3.3 Functional connectivity

#### 3.3.1 Visual seeds in the early blind

In order to investigate the resting-state functional connectivity (rsFC) of the visual areas activated by our 4 cognitive paradigms, we used their respective contrasts of interest to define seed regions. For each of the “Semantics”, “Verb generation”, “Executive Control” and “Long-term Memory” contrasts, one seed was defined using all the voxels activated by the contrast in the visual cortex of blind subjects (Fig. 2). One may note that the activation maps overlap somewhat between seeds. In principle, such overlap could result from the spatial blurring due to averaging across subjects.

However, using a permutation analysis comparing within-subject to between-subject overlap for all contrast pairs, we found that those overlapping activations could not be reduced to cross-subjects averaging (Supplementary Materials and Methods; Supplementary Table 1; Supplementary Fig. 4).

#### 3.3.2 Whole-brain connectivity

We subsequently used the four seeds in an rsFC analysis on resting-state data collected in the same subjects. We first computed the rsFC from those seeds using Pearson’s correlation coefficient for each group separately (Fig. 3), and then compared blind and sighted subjects (Fig. 4). Overall, the 4 seed regions were more strongly connected to the auditory and somatosensory cortices in the sighted. Moreover, inter-hemispheric occipital connectivity was stronger in the sighted for the Verb generation, Executive Control and Semantics seeds. In blind subjects, all occipital seeds had stronger connectivity to the prefrontal and parietal cortices, while the Verb generation, Executive Control and Semantics seeds had also stronger connections to the temporal cortex.

**Figure 3.**
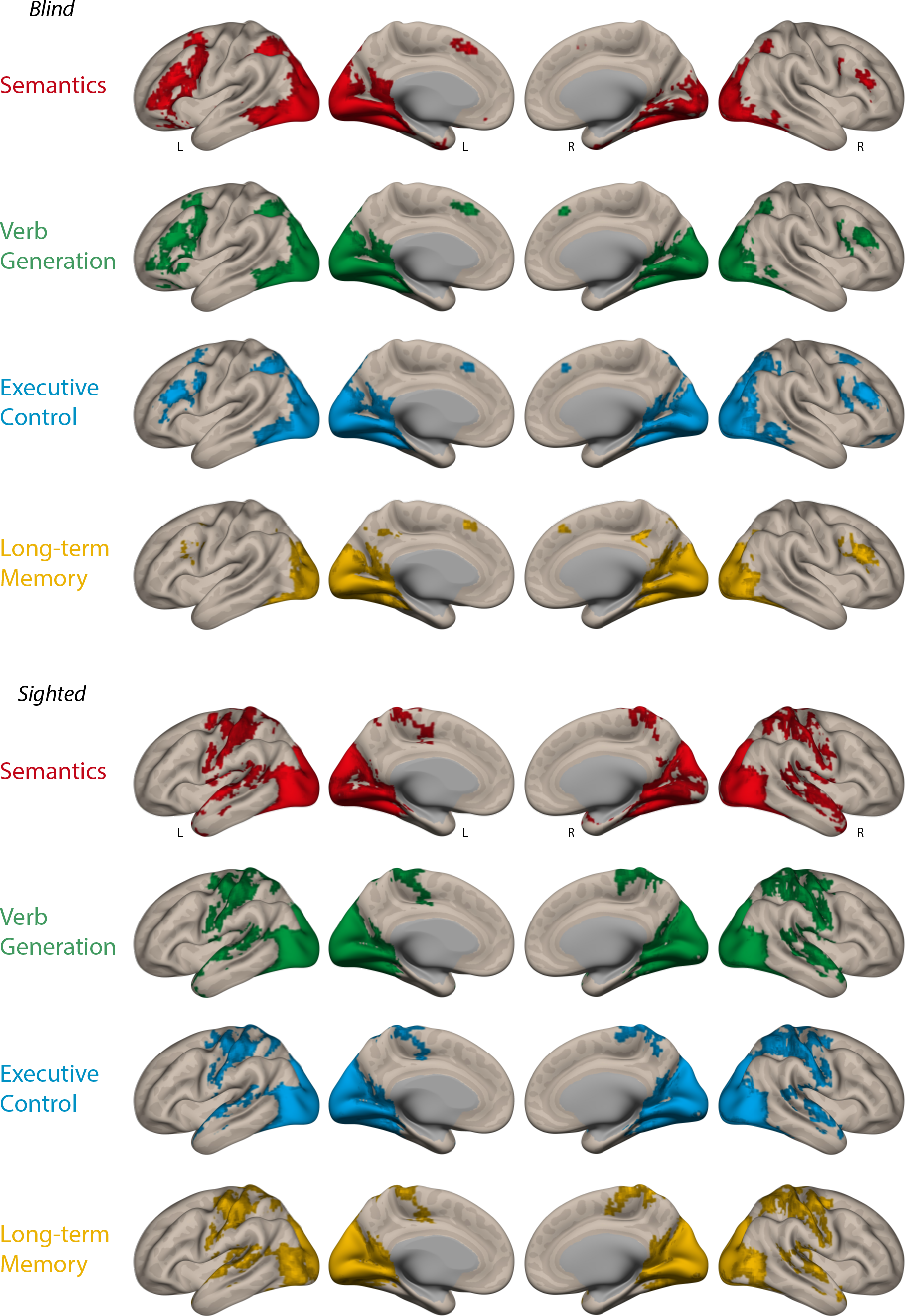
Functional connectivity in the blind and sighted groups separately. Significant functional connectivity seeded from the four occipital regions, using Pearson’s correlation coefficient. Semantics in shown in red, Verb Generation in green, Executive Control in blue, and Long-term Memory in yellow. Voxelwise cluster-forming threshold was set to p<0.005 and all maps are clusterwise corrected (p<0.05).

**Figure 4.**
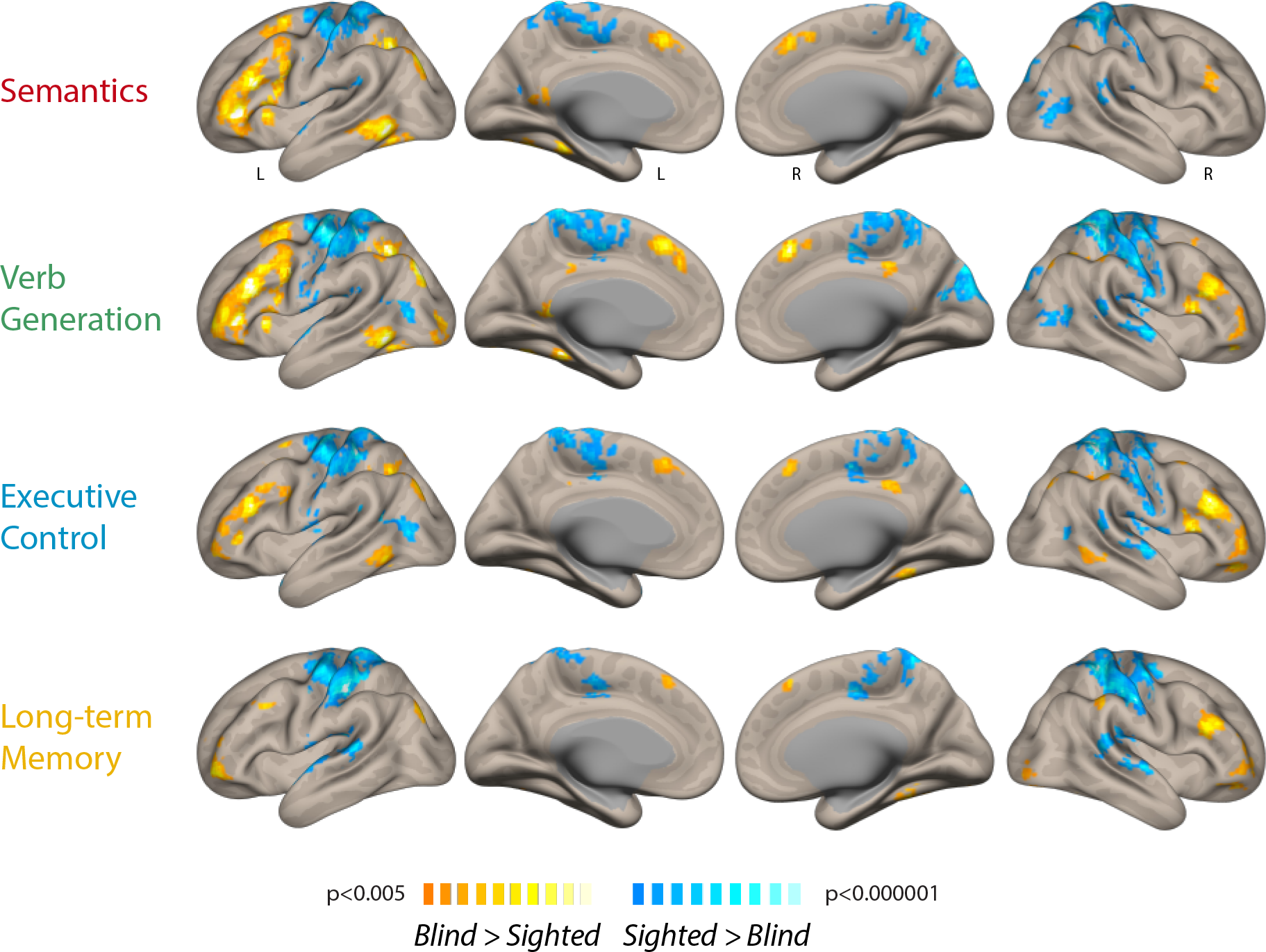
Functional connectivity comparison between the blind and sighted subjects. Significant differences in functional connectivity seeded from the four occipital regions, using Pearson’s correlation coefficient. Hot colors show blind minus sighted differences, and cold colors show sighted minus blind differences. Voxelwise cluster-forming threshold was set to p<0.005 and all maps are clusterwise corrected (p<0.05).

In order to examine the unique connectivity of each seed region, i.e., that which is based only on non-shared signal, we used semi-partial correlations (see Materials and Methods; Fig. 5; Supplementary Fig. 5). In the group of blind subjects, the Verb generation seed was uniquely connected to the left middle and inferior frontal gyri corresponding to premotor cortex and Broca’s area. The Semantics seed was uniquely connected to the left IFG just dorsal to the Verb generation cluster, and to the left lateral temporal cortex, from the occipitotemporal junction to the MTG/STS and the temporal pole. The Executive Control seed was uniquely connected to the right fronto-parietal executive network, in addition to the left IPS and right precuneus. The Long-term Memory seed was uniquely connected to a region in the right precuneus and the bilateral anterior cingulate cortex (ACC). A similar analysis, performed on smaller non-overlapping seeds equated in size, showed similar results (Supplementary material; Supplementary Fig. 6).

**Figure 5.**
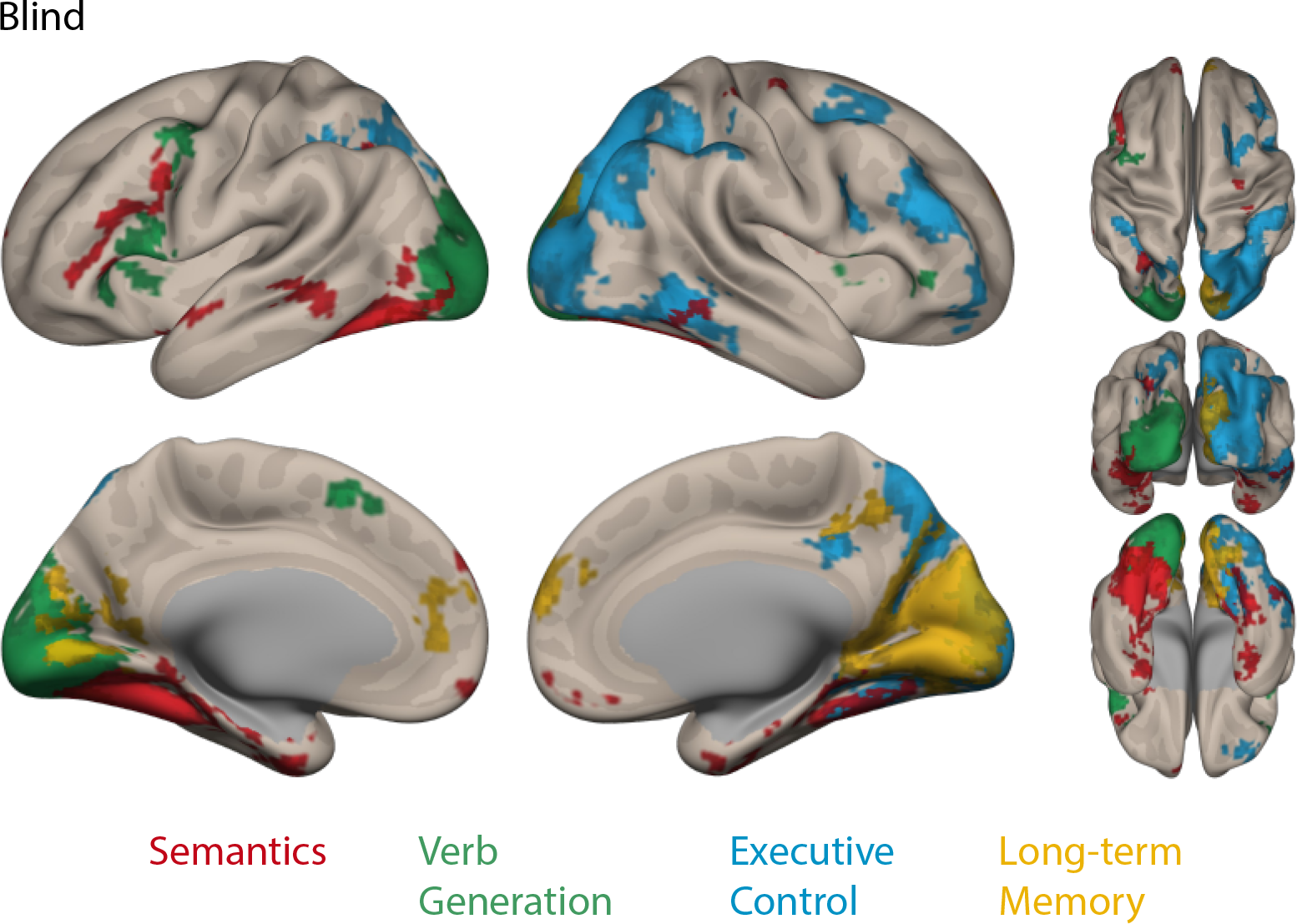
Unique connectivity fingerprints of the occipital seeds in blind subjects. Functional connectivity from the four occipital seeds as delineated using semi-partial correlation. Semantics in shown in red, Verb Generation in green, Executive Control in blue, and Long-term Memory in yellow. Voxelwise cluster-forming threshold was set to p<0.005 and all maps are clusterwise corrected (p<0.05). See Supplementary Fig. 5 with the different maps displayed on separate surfaces, Supplementary Fig. 6 for seeds equated on size, and Supplementary Fig. 8 for an identical analysis in sighted subjects.

We predicted that those regions uniquely connected to visual seeds in the blind should be sensitive to the contrasts that were used to define the corresponding seeds. To answer this question, we defined ROIs using the unique connectivity maps at the individual level (see Materials and Methods). We found that indeed all four contrasts were significant in the ROIs defined by their unique connectivity patterns, with the exception of the semantic contrast in the left prefrontal ROI (Table 3; See also Supplementary Table 2 for results of all contrasts in all ROIs).

**Table 3.** Task-related activation in uniquely connected regions.

|  | Region | Contrast | t-statistic | FDR-corrected p-value |
| --- | --- | --- | --- | --- |
| **a** | Left superior/middle temporal | Semantics | 2.654 | p=0.034 |
| **b** | Left prefrontal | Semantics | 1.744 | p=0.119 |
| **c** | Left prefrontal | Verb generation | 2.811 | p=0.032 |
| **d** | Right prefrontal | Executive control | 2.900 | p=0.032 |
| **e** | Right superior parietal | Executive control | 3.424 | p=0.021 |
| **f** | Anterior cingulate cortex | Long-term memory | 3.879 | p=0.021 |
| **g** | Precuneus | Long-term memory | 3.664 | p=0.021 |

In the same ROIs, we computed the percent variance explained uniquely by each of the four seeds (semi-partial correlation), and the variance explained when combining them together (Table 4). We found that the variance explained by the seeds used to define those ROIs was always larger than the variance explained by any other seed (Table 4, values in bold). This uniquely explained variance amounted to about half the variance explained by a full regression model incorporating all seeds.

**Table 4.**
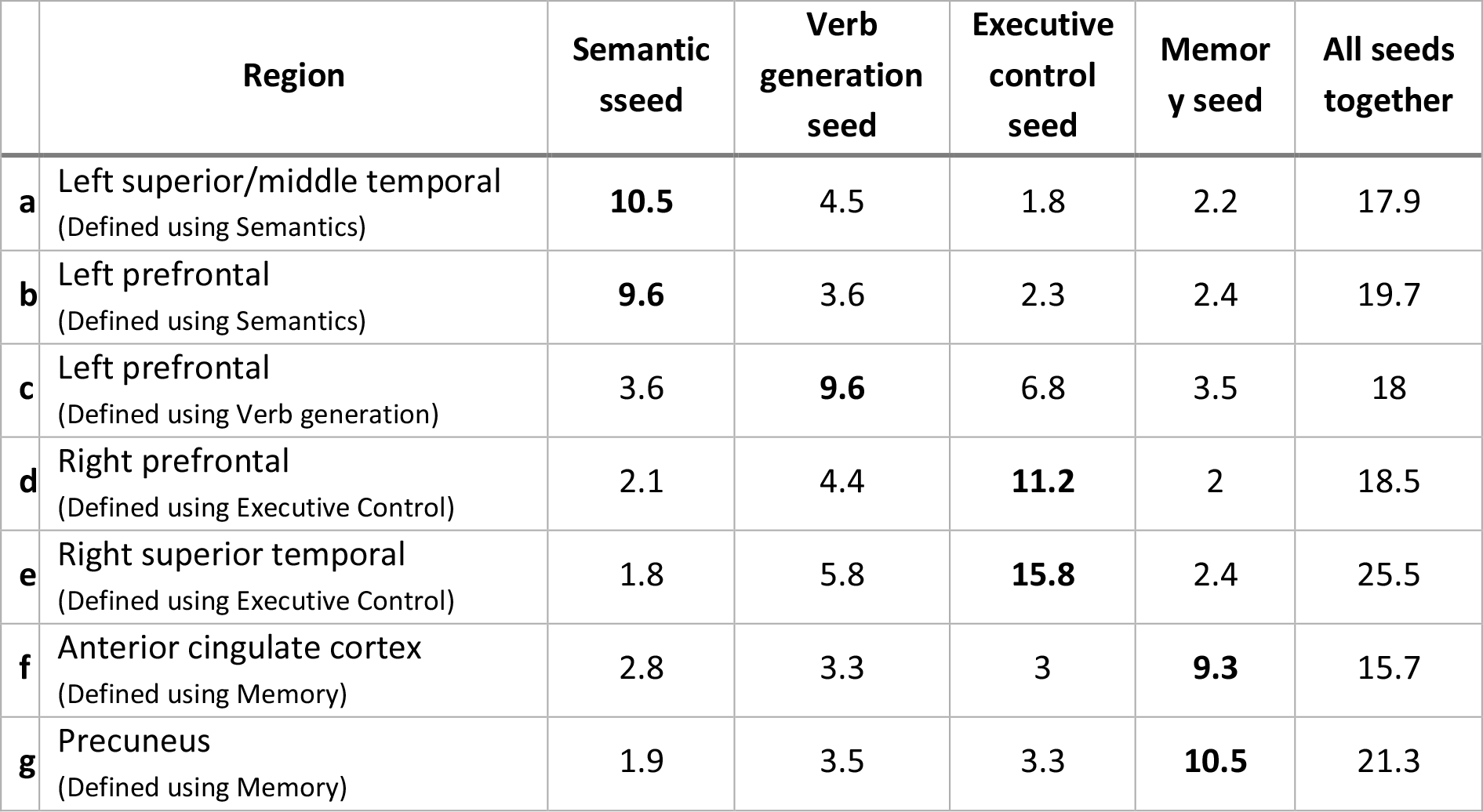
Percent variance in uniquely connected regions explained by semi-partial correlations.

|  | Region | Semantic<br>sseed | Verb<br>generation<br>seed | Executive<br>control<br>seed | Memor<br>y seed | All seeds<br>together |
| --- | --- | --- | --- | --- | --- | --- |
| **a** | Left superior/middle temporal<br>(Defined using Semantics) | **10.5** | 4.5 | 1.8 | 2.2 | 17.9 |
| **b** | Left prefrontal<br>(Defined using Semantics) | **9.6** | 3.6 | 2.3 | 2.4 | 19.7 |
| **c** | Left prefrontal<br>(Defined using Verb generation) | 3.6 | **9.6** | 6.8 | 3.5 | 18 |
| **d** | Right prefrontal<br>(Defined using Executive Control) | 2.1 | 4.4 | **11.2** | 2 | 18.5 |
| **e** | Right superior temporal<br>(Defined using Executive Control) | 1.8 | 5.8 | **15.8** | 2.4 | 25.5 |
| **f** | Anterior cingulate cortex<br>(Defined using Memory) | 2.8 | 3.3 | 3 | **9.3** | 15.7 |
| **g** | Precuneus<br>(Defined using Memory) | 1.9 | 3.5 | 3.3 | **10.5** | 21.3 |

This implies that the unique connectivity reported in this study accounts for a large portion of the explained variance and is not merely a thin layer of modulation in those networks.

Furthermore, as measurement noise could potentially affect semi-partial correlations (Liu 1988), we compared connectivity patterns using a second approach. We performed pair-wise comparisons between the maps of Pearson’s correlation associated to the four seeds (see Supplementary Materials). The maps resulting from those comparisons essentially replicated the results found using semi-partial correlations (Supplementary Fig. 7). For example, the Verb generation seed was more connected than the Executive control seed to the left IFG, while in the opposite direction connections prevailed to the right fronto-parietal executive network (Supplementary Fig. 7a). Moreover, the Semantics seed was more connected to the left MTG/STS and IFG when compared to the Executive control seed, which was more connected to fronto-parietal network bilaterally (Supplementary Fig. 7b). Finally, the Long-term memory seed was more connected to the right precuneus when compared to the Verb generation seed, which was more connected to the left DLPFC (Supplementary Fig. 7e).

For completeness, we also provide the results of the unique connectivity analysis in the sighted. As expected, unique connectivity is limited to regions which have stronger correlation to the visual cortex than in the blind, that is, bilateral auditory and somatosensory cortices (Supplementary Fig. 8).

## 4 Discussion

Using a combination of task and resting-state fMRI, we tested two predictions about the involvement of the visual cortex of early blind individuals in high-level cognitive functions. Each prediction will be discussed in turn, followed by addressing the links between our results and theories of brain reorganization under sensory deprivation. Unless stated otherwise, visual cortex will refer to the visual cortex of blind individuals.

Our first prediction was that high-level cognitive tasks should elicit activations in the visual cortex of blind subjects relative to sighted controls.

### 4.1 Non-perceptual activations in the visual cortex

All experiments showed that, relative to rest, non-perceptual tasks elicited activations in the visual cortex of the blind vs. sighted subjects. Moreover, tighter contrasts controlling for low-level parameters demonstrated that those activations were actually related to high-level cognitive processes.

#### 4.1.1 Speech perception

Our results agree with previous demonstrations that various aspects of language processing activate the visual cortex of blind individuals, including Braille reading (Sadato et al. 1996; Cohen et al. 1997; Beisteiner et al. 2015), word generation (Burton et al. 2002; Amedi et al. 2003; Watkins et al. 2012), speech perception (Bedny et al. 2011, 2015), semantic retrieval (Noppeney et al. 2003) and sentence-level syntax and semantics (Röder et al. 2002; Lane et al. 2015). Using a design similar to ours, Bedny et al. (2011) found sensitivity to semantics, but also to syntax, for which we only found sub-threshold activation. When they compared blind to sighted subjects using a semantics contrast, they found significant differences that peaked in the posterior fusiform gyrus at a distance of 18 mm from the peak of the equivalent analysis in our study. However, they also found stronger activation in the left middle and inferior occipital gyri. Sensitivity to syntax, as well as more extensive activations, may result from their use of a demanding working-memory task, while we chose a shallower perceptual task, in order to equate attentional engagement and difficulty across conditions.

#### 4.1.2 Word generation

The word generation task involves both language and executive functions and activates a large part of the visual cortex with strong left predominance in similarity to results using auditory or Braille words (Burton et al. 2002; Amedi et al. 2003; Ofan and Zohary 2007; Struiksma et al. 2011). However, in a similar task or auditory naming, anophthalmic subjects activated only the lateral occipital cortex bilaterally when compared with controls (Watkins et al. 2012). This more restricted bilateral activation could be related to the aetiology of blindness or to the small group size. In our study, as the task included a verbal semantic component, activations included left ventral cortex, similar to the Semantic contrast. Moreover, they extended to left-predominant lateral occipital areas which may be involved in the executive component. Indeed, those left lateral areas were roughly symmetrical to those activated in the right hemisphere by the non-verbal executive task, to be discussed next.

#### 4.1.3 Executive control

Occipital activation by blocks requiring task-switching indicates involvement of the visual cortex in executive task-set reconfiguration (Monsell 2003), typically a dorsolateral prefrontal function (Koechlin et al. 2003). This novel finding fits with the increased resting-state connectivity observed in blind individuals between visual and prefrontal regions (Liu et al. 2007; Burton et al. 2014). Previous studies of executive control in blind individuals targeted working memory. Using tactile stimulation, Burton et al. (2010) found no evidence of the visual cortex contributing to working memory (Burton et al. 2010). However, (Deen et al. 2015) combining fMRI activation and functional connectivity provided indirect evidence that the visual cortex is involved in auditory working memory. Note that a wide variety of tasks recruit executive processes, and due to the lack of stringent control conditions, some of the visual activations observed in early studies of language and memory (Sadato et al. 1996; Burton et al. 2002; Amedi et al. 2003) may be accounted for by executive cost (Lewis et al. 2010; Park et al. 2011). To the best of our knowledge, this is the first study directly showing the recruitment of non-verbal executive control in blind individuals. This does not rule out, however, that the right lateral visual cortex might intervene in other cognitive functions such as spatial attention and numerical processing (Collignon et al. 2011; Kanjlia et al. 2016).

Considering the additional involvement of the occipital cortex in executive control, one might expect better task performance in blind subjects. However, the behavioral results of this experiment show that blind subjects performed actually worse than controls in the Dual-task condition. In general, the behavioral literature of executive functions in blind subjects is very heterogeneous. Several studies show an executive advantage in blind subjects (e.g. Crollen et al. 2011; Withagen et al. 2013 in blind children; Dormal et al. 2016) while others find no differences between blind and sighted subjects (e.g. Bliss et al. 2004; Wan et al. 2010; Pigeon and Marin-Lamellet 2015). Executive abilities in the blind were also shown to depend on task complexity and perceptual stimulus parameters (e.g. Rokem and Ahissar 2009; Occelli et al. 2017). Concerning the contribution of the visual cortex to executive processing, it is not at all clear that a more extended activation or a bigger network should necessarily lead to a better performance.

#### 4.1.4 Long-term memory

The mesial visual cortex was activated during successful retrieval in an incidental long-term memory task (i.e. for hits over correct rejections). Those activations overlap with the bilateral activations observed when EB recall a list of abstract words (Amedi et al. 2003) or perform an old/new judgment (Raz et al. 2005). Both studies support a causal involvement of the visual cortex in memory retrieval by showing a correlation between activation level in the Calcarine sulcus and individual performance. Consistent with these findings, the peak difference between blind and sighted subjects in our memory contrast also falls within the limits of the Calcarine sulcus (see Results). Using both auditory and Braille words, Burton et al. (2012) showed that such memory-related activations occur irrespective of input modality, but with no advantage for successful retrieval (Burton et al. 2012).

Overall difficulty was equated across the compared conditions in the long-term memory and speech perception experiments using carefully matched sentences and an orthogonal task. Hence, activations cannot be accounted for by non-specific effects of difficulty. However, in the executive control and the word generation experiments, conditions do differ in terms of task difficulty. For those tasks, this confound should not be taken as a shortcoming, as task difficulty is largely synonymous to executive toll, which is precisely the function we wish to assess (Gilbert et al. 2012).

Having discussed the four tasks separately, we now consider the occipital voxels that were found to be activated by more than one contrast (Fig. 2). Most notable is the overlap between the Verb Generation and the other three contrasts, which may arise from overlapping cognitive functions.

Indeed, the Verb minus Repeat contrast encompasses cognitive processes related to semantic access, retrieval from memory and executive control. By comparing within-subject vs between-subject overlap, we showed that this overlap cannot result only from cross-subject averaging and blurring, but actually reflect the intervention of some voxels in multiple cognitive tasks (Supplementary Fig. 4, Supplementary Table 1).

In summary, we found that all tested cognitive processes activate the visual cortex in the blind subjects. Moreover, as we used task-specific high-level controls, we may conclude that visual activations in each experiment reflect the related cognitive processes probed by each experiment, e.g., memory, language and executive processing.

We now discuss our second prediction, i.e., that visual areas activated under those paradigms are connected to distinct functional networks reflecting their novel functional role.

### 4.2 Integration in brain-scale functional networks

In agreement with previous studies, we found that, in blind subjects, visual seeds generally showed (1) a reduced inter-hemispheric connectivity with contralateral regions; (2) a reduced connectivity with auditory, somatosensory and motor areas; and (3) an increased connectivity with associative cortex in the lateral prefrontal, superior parietal, and mid-temporal areas (Liu et al. 2007; Yu et al. 2008; Bedny et al. 2011; Watkins et al. 2012; Butt et al. 2013; Qin et al. 2013; Burton et al. 2014; Wang et al. 2014; Deen et al. 2015; see however Heine et al. 2015).

The decreased connectivity with auditory and somatosensory areas, which has been observed repeatedly, may still come as a surprise when considering the activation of the visual cortex by auditory and tactile tasks in blind subjects (e.g. Sadato et al. 1996; Collignon et al. 2011). However, Pelland et al. (2017) demonstrated that during an auditory task, unlike during rest, there is an increase in functional connectivity between occipital and temporal regions in blind subjects. This means that caution is required when interpreting such decreases in rsFC.

Beyond commonalities between connectivity maps from the four seeds, we were mostly interested in the unique connectivity of each seed region, which we isolated using semi-partial correlation.

Confirming our prediction, visual seed regions showed unique connectivity with distant task-related networks. First, language-related seeds, derived from the Semantics and Verb generation contrasts, were connected to core language areas, i.e., Broca’s area and the left lateral and anterior temporal lobe, both consistently involved in verbal semantics (Patterson et al. 2007; Binder and Desai 2011). Moreover, we found that those areas are activated by the Semantics and Verb generation contrasts in individual subjects (Table 3). In agreement with this result, Bedny et al. (2011) using seed-based functional connectivity and (Watkins et al. 2012) using ICA decomposition showed visual cortex connectivity with parts of the language network. Moreover, Kanjlia et al. (2016) showed that this fronto-occipital connectivity is specific to language over mathematics. Second, the non-verbal Executive Control seed was mainly connected to a right-hemispheric frontoparietal executive network. In general, task-switching activates bilateral frontoparietal areas (Kim et al. 2012) but we also found that the executive control contrast significantly activates connected regions in this network (Table 3). The present right-lateralization may result from the use of non-verbal material (Yeung et al. 2006; Geddes et al. 2014) but also from the reduced interhemispheric correlation in the visual cortex of blind individuals. The decoupling of the left and right visual cortices may reflect their recycling by different cognitive functions, implying distinct and asymmetric connectivity patterns. In agreement with this hypothesis, in the sighted, the connectivity of three out of the four visual seeds was symmetrical (Supplementary Fig. 8). Last, the Long-term Memory seed was uniquely connected to the precuneus and the bilateral medial prefrontal cortex, which are consistently involved in episodic memory retrieval, particularly in a contrast of recollection vs. familiarity (Spaniol et al. 2009; Kim 2010). Similarly, we also found that those regions are activated by the Long-term memory contrast in blind subjects (Table 3). Note that the use of semi-partial correlation isolates only what is uniquely connected to each seed. Therefore, partial overlap between seed regions is considered as shared signal that is regressed out from both seeds. Any connectivity contributed by those overlapping regions is invisible to this analysis.

It has been previously proposed that the increased connectivity between frontal and occipital regions in blind individuals reflects increased gating of occipital responses by frontal cortex that is not function-specific (Bock and Fine 2014). However, later studies showed that visual regions activated by language and mathematics are preferentially connected to long-distance regions processing language and mathematics, respectively (Kanjlia et al. 2016). Our results support a function-specific account of occipital rsFC in blind individuals by showing functional correspondence between activations and connected networks for a large set of cognitive functions. Those include non-verbal executive processes, which were never directly tested in blind individuals before.

Before moving on to the general discussion, we would like to discuss two methodological points. First, the term connectivity should be employed with caution in the present context. The resting-state functional connectivity (rsFC) results reported here are based on correlation measures between different regions in the brain at rest. Those measures are correlated with anatomical connectivity, but also go beyond it, featuring functional connections between regions that are not directly connected structurally (Honey et al. 2009). Moreover, those measures were shown to be open to contamination by fluctuations originating from non-neuronal sources such as respiration and stomach activity (Birn et al. 2008; Rebollo et al. 2018). They could also be sensitive to the alterations in neural and vascular structures found in blind individuals (Wanet-Defalque et al. 1988; Veraart et al. 1990; Uhl et al. 1993; De Volder et al. 1997; Weaver et al. 2013). However, using a combination of rsFC and task-based activation we were able to show that functionally connected long-distance regions are also activated by the same task. Those congruent findings make it unlikely that the connectivity findings arise only due to non-neuronal factors. Second, it is fair to mention that the use of a volumetric template in MNI space could be a limiting factor in our study. Due to cross-subject variability in the folding patterns of the cortical surface (Zilles et al. 1997), one risks averaging across functionally different regions when aligning individuals to a volumetric template (Hellier et al. 2003). This limitation in spatial accuracy could potentially decrease the power of peak-level inference and increase the power of cluster-level inference (Tucholka et al. 2012). Therefore, the use of surface-based alignment techniques might be preferred for future studies (e.g. Desai et al. 2005).

### 4.3 General discussion

How do our results relate to general accounts of cortical plasticity in blind individuals? Beyond cross-modal plasticity, a generic term to describe the activation of sensory-deprived cortex by the remaining intact senses (Rauschecker 1995; Renier et al. 2014), the activation of the visual cortex by non-visual stimuli may correspond to various perceptual (e.g. motion) and non-perceptual (e.g. memory) processes. According to the metamodal/supramodal theory, the cortex is organized into task operators that are indifferent to sensory modality (Pascual-Leone and Hamilton 2001; Pietrini et al. 2004). Hence, in the case of visual deprivation, auditory and tactile input to the visual cortex would be unmasked, leading to its involvement in non-visual perception (Cecchetti et al. 2016). With the prediction that the same computations are carried out even in blindness, this theory cannot explain the drastic change from low-level visual processing to high-level non-perceptual computations.

It has been recently hypothesized that such reorganization of deafferented visual cortex is driven and shaped during development by pre-existing anatomical connections to frontal, parietal and temporal areas (Bedny 2017). Our results are in line with such a connectivity-based account, as illustrated most clearly in the case of the left ventral language-related seeds (Semantics and Generation), which overlap with the Visual Word Form Area (VWFA). This region is involved in written words recognition in literate subjects, and has privileged anatomical connections to language areas in literate adults (Bouhali et al. 2014), but also in children before reading acquisition (Saygin et al. 2016). Similarly, the Executive Control and Generation seeds extend into lateral occipital cortex, which has direct anatomical connections to prefrontal regions through the IFOF and SLF (Forkel et al. 2014). Finally, the cingulum may connect the occipital Long-term Memory seed to mesial frontal and parietal regions with which it establishes novel functional links in blind individuals (Catani et al. 2002), although the existence of occipital cingulum fibers is still disputed (Rojkova et al. 2016).

How does this connectivity-based account of brain organization reconcile with evidence for a task-based metamodal brain organization? A plausible proposal argues that regions only seem to show functional constancy but in reality their underlying computations are different in blindness (Bedny 2017). For example, a connectivity-based account would claim that motion selectivity in a certain region is retained in blindness because of the motion-relevant information it receives. In this specific case, the two theories diverge on the nature of the computation which is carried out in sighted and blind individuals, requiring further research.

We found activations for high-order functions that overlap with regions that were argued to be metamodal. For example, the left occipital-temporal activation by the semantics contrast, using a high-level auditory language task, overlaps with the VWFA, which has been proposed to be metamodal and activated by reading in both sighted and blind individuals (Reich et al. 2012; Striem-Amit et al. 2012). Some studies suggest that the VWFA requires visual experience to be activated by reading, and the lack thereof would recruit it for high-order language processing (Kim et al. 2017).

Other studies, however, provided evidence in support of VWFA activation by letters and words but not by a subsequent semantic task (Sigalov et al. 2016). Studies targeting the structure of the information processed in this region might contribute to a better understanding of its computation in blindness.

Finally, another open question concerns the relative contribution of, on the one hand, occipital regions involved in a given cognitive process in blind subjects only, and, on the other hand, classical regions operating both in blind and sighted individuals. For example, in the Semantics contrast, ventral occipitotemporal activations did not replace lateral temporal regions, but appear rather as an add-on (Fig. 1a). In the case of auditory motion perception by early blind subjects, it has been shown that direction discrimination is enhanced in the blind-specific hMT+, and reduced in the planum temporale, pointing to a shift in functionality even in classical areas (Jiang et al. 2014). The experimental designs we used do not allow us to test such hypothesis. Moreover, the issue of role distribution between occipital and classical areas should take into consideration inter-subject variability. In different subjects, processing may be balanced differently across the two sets of regions. In the example of language, one would then expect, that subjects with more lateral temporal activation would show less occipital activation, and vice-versa. However, we found that in most cases, long-distance regions connected to a seed in the visual cortex were sensitive to the same contrasts as this seed, suggestive of functional commonality.

In conclusion, using a broad set of language, memory, and executive tasks within the same blind subjects, we demonstrate the selective integration of different patches of the visual cortex into brain-scale networks with distinct topography, lateralization, and functional roles. Identifying the informational content and causal role of those occipital activations will be the main future challenge.

## Acknowledgments

The authors would like to thank A. Marchand for coordinating the data acquisition and S. Mohand-Said for the help in recruiting subjects.

